# Redetermination of the first unknown protein MicroED structure by high resolution X-ray diffraction

**DOI:** 10.1101/2021.04.07.438860

**Authors:** Hongyi Xu, Xiaodong Zou, Martin Högbom, Hugo Lebrette

## Abstract

Microcrystal electron diffraction (MicroED) has the potential to considerably impact the field of structural biology. Indeed, the method can solve atomic structures of a wide range of molecules, beyond the reach of single particle cryo-electron microscopy, exploiting crystals too small for X-ray diffraction (XRD) even using X-ray free-electron lasers. However, until the first unknown protein structure – a R2-like ligand-binding oxidase from *Sulfolobus acidocaldarius* (*Sa*R2lox) – was recently solved at 3.0 Å resolution, MicroED had only been used to study known protein structures previously obtained by XRD. Here, after adapting sample preparation protocols, the structure of the *Sa*R2lox protein originally solved by MicroED was redetermined by XRD at 2.1 Å resolution. In light of the higher resolution XRD data and taking into account experimental differences of the methods, the quality of the MicroED structure is examined. The analysis demonstrates that MicroED provided an overall accurate model, revealing biologically relevant information specific to *Sa*R2lox, such as the absence of an ether cross-link, but did not allow to detect the presence of a ligand visible by XRD in the protein binding pocket. Furthermore, strengths and weaknesses of MicroED compared to XRD are discussed in the perspective of this real-life protein example. The study provides fundaments to help MicroED become a method of choice for solving novel protein structures.

**Synopsis:** The first unknown protein structure solved by microcrystal electron diffraction (MicroED) was recently published. The redetermination by X-ray diffraction of this protein structure provides new insights into the strengths and weaknesses of the promising MicroED method.

## 1. Introduction

In a near future, microcrystal electron diffraction (MicroED), also referred as 3D electron diffraction (3D ED), could become one of the indispensable methods in the toolbox of structural biologists, routinely available to solve specific challenging problems (Clabbers & Xu, 2021; Gemmi *et al*., 2019; Nannenga & Gonen, 2019). It could complement other main techniques of the field, *i*.*e*., cryo-electron microscopy (cryo-EM) and X-ray diffraction (XRD), in term of sample requirements and molecular targets. Indeed, MicroED uses crystals too small for single-crystal XRD (Nannenga, 2020), or even for serial crystallography at X-ray free-electron laser sources (XFEL) (Wolff *et al*., 2020), and enables the study of small molecules inaccessible to single particle cryo-EM (Jones *et al*., 2018; Kunde & Schmidt, 2019). Therefore, the development of this emerging approach could greatly benefit drug discovery (Brázda *et al*., 2019; Clabbers *et al*., 2020; Clabbers & Xu, 2020). For small molecule crystallography, using 3D ED/MicroED, sub-Ångström resolution data from inorganic and organic crystals are easily obtained, from which accurate atomic structure models can be determined (Huang *et al*., 2021). However, to live up to expectations in structural biology, MicroED still has to prove that it can elucidate unique problems and provide atomic models that meet high-quality standards established by X-ray protein crystallography.

MicroED had only been applied for structure determination of known proteins until we recently solved the first unknown structure of a new protein, the R2-like ligand-binding oxidase from *Sulfolobus acidocaldarius* (*Sa*R2lox) (Xu *et al*., 2019). The R2lox metalloenzyme family share a common fold with the ribonucleotide reductase R2 protein but its physiological function is still unknown. Two crystal structures of R2lox proteins had been solved previously by XRD, from *Mycobacterium tuberculosis* and *Geobacillus kaustophilus* (PDB ID: 3ee4 and 4hr0 for *Mt*R2lox and *Gk*R2lox, respectively) (Andersson & Hogbom, 2009; Griese *et al*., 2013). In both cases, structures revealed an α-helix bundle core accommodating a Mn^III^Fe^III^ dinuclear metal cluster interacting with a long-chain fatty acid, and a unique feature: a tyrosine-valine ether cross-link between the two residue side-chains. One of these structures was used as a search model for molecular replacement (MR) to phase the MicroED data that allowed us to solve the novel *Sa*R2lox structure at 3.0 Å resolution (PDB ID: 6qrz). In order to produce the best atomic coordinates despite the limited resolution, we merged data from 21 crystals, as we previously demonstrated that merging MicroED data from a large set of crystals significantly improves not only the data completeness and overall signal-to-noise ratio but also the quality of the final model (Xu *et al*., 2018). The structure provided new biological insights on this family of proteins, exhibiting a distinct substrate-binding pocket with specific surrounding residues and electrostatic contact potentials. The quality of the electrostatic potential map allowed us to model a dinuclear metal centre but neither a ligand nor the tyrosine-valine ether cross-link, bringing into question their presence in this protein. However, in absence of a high-resolution structure solved by a well-established technique of structural biology, it was difficult to evaluate the quality of the final model in its entirety.

Here we present the structure of the same *Sa*R2lox protein redetermined using XRD. By modifying the crystallization protocol in order to meet the requirements of single-crystal XRD, we were able to obtain a structure at 2.1 Å resolution. Data reduction and model refinement were performed in a similar way as for structure solved by MicroED. In addition, different MR and refinement approaches were tested. Our results allow us to proceed to an in-depth assessment of the MicroED data and resulting atomic coordinates. We show that the MicroED data allowed accurate modelling of most of the protein structure. However, despite unveiling several biologically relevant features specific to *Sa*R2lox, such as the absence of the ether cross-link or the correct metal–metal distance of the cofactor, MicroED failed at detecting the presence of the ligand as revealed by XRD. Finally, we discuss the challenges encountered to determine the *Sa*R2lox structures by both MicroED and XRD in order to highlight strengths and weaknesses of MicroED and pave the way for possible improvements.

## 2. Methods

### 2.1. Crystallisation and sample handling

Protein R2-like ligand-binding oxidase from *Sulfolobus acidocaldarius* (*Sa*R2lox) used in this study belongs to the same batch used to solve the structure by MicroED using a transmission electron microscope (TEM); production and purification procedures are described in (Xu *et al*., 2019). *Sa*R2lox was crystallized using the hanging drop vapor diffusion method. The final condition consisted in a volume of 1 μl of protein solution at 8 mg/ml mixed with 0.5 μl of reservoir solution made of 43% (v/v) polyethylene glycol (PEG) 400, 0.2 M lithium sulphate and 0.1 M sodium acetate pH 4.5. Within 1 to 2 days at 21°C, clusters of thin platelike crystals stacked on top of each other and measuring individually around 100 x 100 x 10 µm^3^ appeared in the drop.

After opening the well, single crystals were manually detached from clusters and harvested using cryo-loops in less than one minute. Opening the well resulted in drops becoming rapidly opaque rendering impossible to distinguish the crystals from the mother liquor, even using polarized light. Then crystals were flash-cooled in liquid nitrogen without additional cryo-protecting agent.

### 2.2. Data collection and data processing

XRD data were collected on the beamline X06SA (PXI) at the Swiss Light Source (SLS), equipped with an EIGER X 16M (Dectris) detector at a wavelength of 1.0 Å at cryogenic temperature. Fine slices of 0.1° were collected over a total range of 360°. Data reduction was performed using *XDS* (Kabsch, 2010), initially without inputs on the space group or unit cell. Bravais lattice and unit cell parameters were automatically determined by *XDS*. Additionally, the two screw axes were inferred from the presence of the systematic absences. Unless stated otherwise, reflections h, k, l and -h, -k, -l were treated as equivalent, considering the Friedel’s law true in *XDS*, similarly as for the MicroED data processing. The reflection file was subsequently converted to mtz format using *XDSCONV*. The high-resolution cut-off was determined based on a combination of I/σ(I), *R*_*meas*_ and CC_1/2_. Data reduction statistics were obtained from (Xu *et al*., 2019) for MicroED and compiled using *phenix*.*table_one* for XRD.

The *R*_*free*_ test set defined in the MicroED data was conserved for the XRD dataset by importing the *R*_*free*_ test set flag (FreeR_flag) and extending to resolution and completeness to represent approximately 5% of total reflections using *phenix*.*reflection_file_editor*. Structure factor data quality was assessed with *phenix*.*xtriage*. The structure factor file for the MicroED dataset (PDB ID: 6qrz) was obtained in mtz format using the SF-TOOL web server (http://sf-tool.wwpdb.org).

### 2.3. Structure solution and refinement

Phasing was performed by MR using *Phaser* (McCoy *et al*., 2007). The final XRD model deposited in the Protein Data Bank (PDB ID: xxxx) was obtained with a MR search model built from the atomic coordinates of *Mt*R2lox (PDB ID: 3ee4), similarly as for the MicroED structure. Additional MR search models were also tested: *Gk*R2lox solved by XRD, and *Sa*R2lox solved by MicroED (PDB ID: 4hr0 and 6qrz, respectively). In all three cases, a well-contrasted solution was obtained with one molecule per asymmetric unit in the space group *P*2_1_2_1_2. Search models based on *Mt*R2lox and *Gk*R2lox were edited with *Sculptor* (Bunkóczi & Read, 2011).

All subsequent refinement steps were conducted independently from the MicroED model. Likewise the MicroED structure, refinement was performed with *phenix*.*refine* (Afonine *et al*., 2012) using merged intensities, and selecting automatic optimisation of target weights for stereochemistry and *B* factors, but using the default n-gaussian X-ray scattering table instead of the electron scattering table. The refinement strategy started with a rigid body (RB) refinement step followed by refinement cycles of atomic coordinates and individual isotropic *B* factors. Model examination, real-space refinement and manual modifications were iteratively conducted with *Coot* (Emsley *et al*., 2010). When all residues visible in the electron density map were built, and before adding metal ions, ligand or water molecules, *B* factors were reset to 20 and the coordinates were subjected to Cartesian simulated annealing in order to remove any possible model bias. Subsequently, metal ions, waters and a palmitic acid ligand were added, and the model was further refined. Restraints for metal coordination were generated using elbow (Moriarty *et al*., 2009), and manually modified based on previous studies on R2lox structures (Andersson & Hogbom, 2009; Griese *et al*., 2013). Unlike other R2lox proteins metal-loaded in aerobic conditions, no electron density could be observed for the ether cross-link between Tyr174 and Val82, and consequently no corresponding restraints were applied during refinement. Quality of models were evaluated using MolProbity (Williams *et al*., 2018). Additional refinement tests were conducted but were limited to a step of RB followed by three refinement cycles of individual isotropic *B* factors, *i*.*e*., no atomic coordinate refinement was performed. Electron or n-gaussian X-ray scattering table was chosen depending on the nature of the diffraction data used during the refinement. Refinement statistics were generated by *phenix*.*refine* or compiled using *phenix*.*table_one*. Note that refinement statistics for the MicroED structure (PDB ID: 6qrz) were recomputed using *phenix*.*refine* for comparability.

Simulated annealing composite omit maps were calculated with sequential 5% fractions of the structure omitted using *phenix*.*composite_omit_map* with the Cartesian annealing method (only observed reflections were used for map calculations, *i*.*e*., no missing *F*_*obs*_ were restored using a weighted *F*_*calc*_). Simulated annealing omit *F*_*obs*_ *– F*_*calc*_ maps were generated using *phenix*.*refine* after Cartesian simulated annealing refinement were performed on atomic models where palmitic acid and metal-coordinating waters were deleted. Anomalous difference maps were computed with *phenix*.*maps* using Friedel pair reflections unmerged to preserve the contribution of anomalous scattering atoms. Normalized mean *B* factors (including all atoms for each residue) were compiled and plotted using *phenix*.*structure_comparison*. All *phenix* programs belong to the *PHENIX* suite version 1.19_4092 (Liebschner *et al*., 2019). Cα deviation values between structures were obtained using the secondary-structure matching tool (Krissinel & Henrick, 2004). Figures were prepared using Inkscape 1.0 and the PyMOL Molecular Graphics System, version 2.4 Schrödinger, LLC.

## 3. Results

### 3.1. Crystallization and sample handling

From the original crystallization condition producing the microcrystals that made it possible to solve the structure by MicroED, we conducted an optimisation to obtain crystals of suitable size and shape for single-crystal XRD data collection at cryo-temperature at synchrotron (**Fig. 1**). Since we aimed to compare structures obtained by the two methods, we kept consistent as many parameters as possible, *e*.*g*., by using the same batch of purified protein, and collecting data at cryo-temperature. In addition, we put effort into minimising changes to the crystallization condition. The key parameters we modified to allow the growth of bigger crystals were an increase of pH from 3.4 to 4.5, a volume ratio of protein to reservoir solutions from 1:1 to 2:1, and a decrease of the drop volume from 4 µl to 1.5 µl (**Table 1**). This new protocol produces platelike crystals of approximately 100 µm in length grown in clusters and stacked on top of each other.

**Figure 1.**
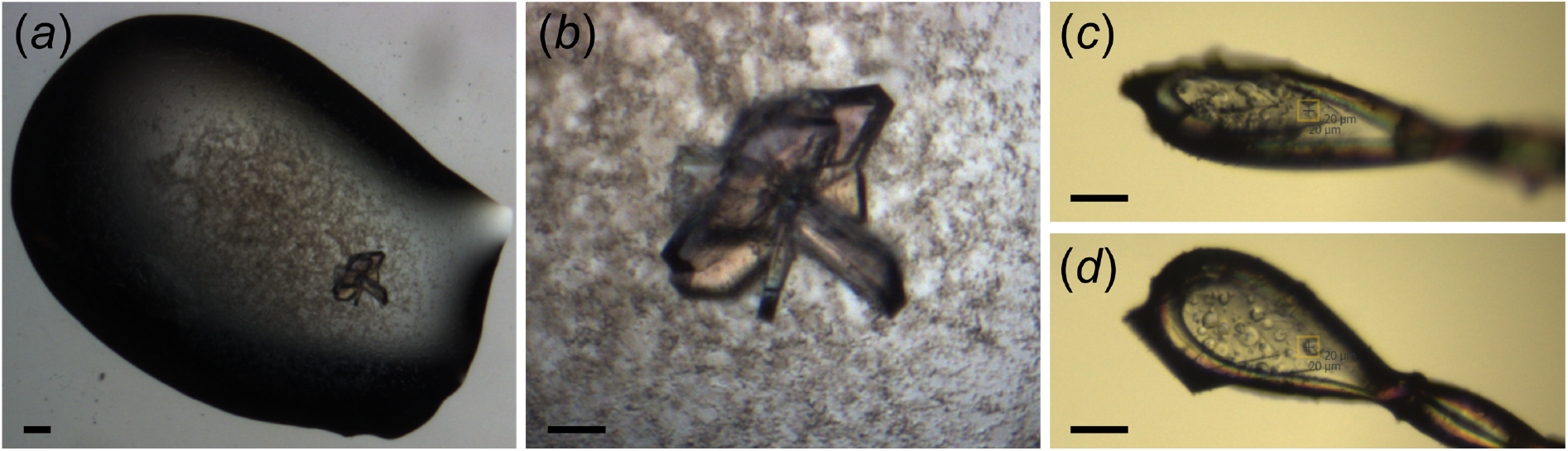
Crystals of *Sa*R2lox optimised for single-crystal XRD data collection. (*a*) Typical *Sa*R2lox crystals used for XRD experiments obtained by hanging drop vapor diffusion and forming a cluster surrounded by precipitate. (*b*) Close-up of the crystals. (*c*) and (*d*) Single crystal mounted in a cryo-loop at the synchrotron beamline pictured at two angles 45° apart. Scale bar 100 µm in (*a*), and 50 µm in (*b*), (*c*) and (*d*).

**Table 1.**
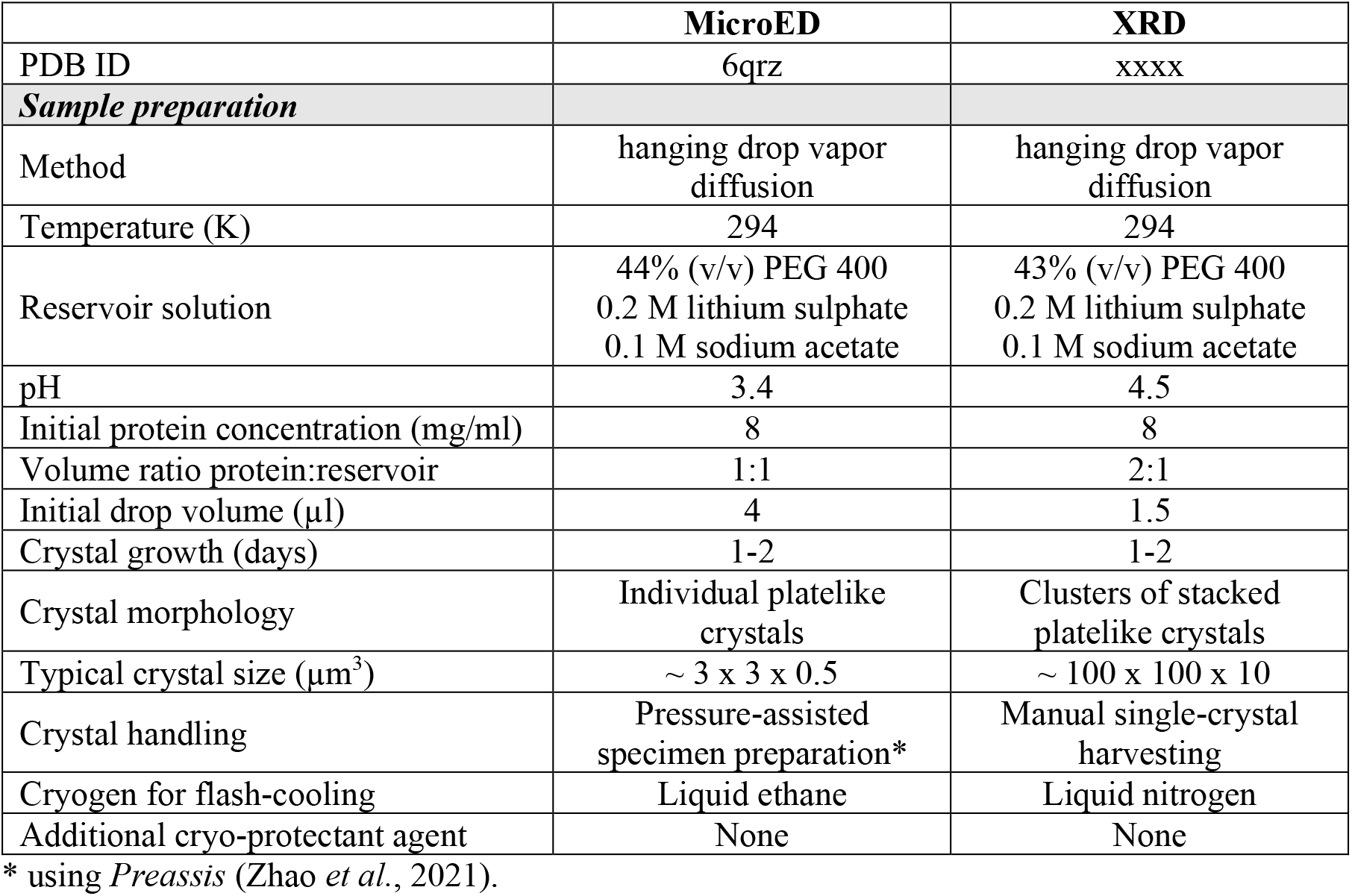
XRD sample preparation of *Sa*R2lox compared to MicroED.

|  | **MicroED** | **XRD** |
| --- | --- | --- |
| PDB ID | 6qrz | xxxx |
| ***Sample preparation*** |  |  |
| Method | hanging drop vapor diffusion | hanging drop vapor diffusion |
| Temperature (K) | 294 | 294 |
| Reservoir solution | 44% (v/v) PEG 400<br>0.2 M lithium sulphate<br>0.1 M sodium acetate | 43% (v/v) PEG 400<br>0.2 M lithium sulphate<br>0.1 M sodium acetate |
| pH | 3.4 | 4.5 |
| Initial protein concentration (mg/ml) | 8 | 8 |
| Volume ratio protein:reservoir | 1:1 | 2:1 |
| Initial drop volume (μl) | 4 | 1.5 |
| Crystal growth (days) | 1-2 | 1-2 |
| Crystal morphology | Individual platelike crystals | Clusters of stacked platelike crystals |
| Typical crystal size (μm<sup>3</sup>) | ~ 3 x 3 x 0.5 | ~ 100 x 100 x 10 |
| Crystal handling | Pressure-assisted specimen preparation* | Manual single-crystal harvesting |
| Cryogen for flash-cooling | Liquid ethane | Liquid nitrogen |
| Additional cryo-protectant agent | None | None |

\* using *Preassis* (Zhao *et al.*, 2021).

Although crystals are of sufficient size, manual harvesting is laborious. Firstly, it is difficult to mount individual crystals because they are thin, fragile and tend to break when separated from each other. Secondly, opening the well breaks the equilibrium and results in drops becoming rapidly opaque. Consequently, crystals have to be fished out within a minute before they become indistinguishable from the rest of the drop. Thirdly, the mother liquor is very viscous due to the high concentration (43% (v/v)) of polyethylene glycol (PEG) 400. These challenges were all avoided in our MicroED experiments as the method does not require handling of single crystals, and because we used pressure-assisted specimen preparation, *Preassis* (Zhao *et al*., 2021), which allows to easily manage very viscous crystallization conditions.

Regarding the diffraction quality, our first observation is that it is not consistent between crystals, probably due to the difficulty in isolating a single crystal without damaging it. Out of 10 crystals from the same optimal crystallization condition, only two crystals displayed good quality diffraction to resolution better than 3 Å. The best dataset we obtained was processed at 2.1 Å resolution, thus notably better than the 3.0 Å resolution obtained using MicroED. To note, the XRD data were collected using a detector EIGER 16M (Dectris) which is currently amongst the best detector to collect noise-free data in macromolecular crystallography (Casanas *et al*., 2016).

### 3.2. Data reduction

XRD data processing was performed similarly as for MicroED, using *XDS* to automatically determine the Bravais lattice and unit cell parameters. The presence of two screw axes could be inferred from the systematic absences in the list of reflections. The resolution of 2.1 Å was defined taking into account I/σ(I), *R*_*meas*_ and CC_1/2_ (**Table 2**). The unit cell dimensions a, b and c are respectively 1.44%, 1.65% and 0.80% longer for XRD data than for MicroED, resulting in an increase of 3.9% in total unit cell volume. The fact that data obtained by MicroED display unusual cell dimensions compared to cryogenic XRD experiments was previously investigated to reach the conclusion that it is not due to measurement error but possibly the effect of experimental conditions (Wolff *et al*., 2020). In our case, we could hypothesize that it results from differences in crystallization pH, or cryo-cooling temperature or pressure (**Tables 1 and 2**).

**Table 2.** XRD data collection and data reduction statistics of *Sa*R2lox compared to MicroED.

|  | **MicroED** | **XRD** |
| --- | --- | --- |
| PDB ID | 6qrz | xxxx |
| ***Data collection*** |  |  |
| Number of crystals | 21 | 1 |
| Crystal size ($\mu\text{m}^3$) | $\sim 3 \times 3 \times 0.5$ | $\sim 100 \times 100 \times 10$ |
| Scattering Type | Electron | X-ray |
| Wavelength ($\text{\AA}$) | 0.0251 | 1.0 |
| Radiation source | TEM JEOL 2100<br>(LaB<sub>6</sub> filament) | Synchrotron SLS<br>beamline X06SA |
| Beam size ($\mu\text{m}^2$) | 3 | 400 |
| Protocol | Continuous rotation | Continuous rotation |
| Detector | Hybrid pixel detector<br>Timepix (Amsterdam<br>Scientific Instruments) | Hybrid pixel detector<br>EIGER X 16M (Dectris) |
| Temperature (K) | 100 | 100 |
| Pressure (Pa) | $10^{-5}$ | $10^5$ |
| ***Data reduction*** |  |  |
| Software | *XDS* | *XDS* |
| Resolution range ($\text{\AA}$) | 29.00-3.00 (3.08-3.00) | 44.47-2.10 (2.175-2.10) |
| Space group | *P2<sub>1</sub>2<sub>1</sub>2* | *P2<sub>1</sub>2<sub>1</sub>2* |
| Unit cell dimensions<br>a, b, c ($\text{\AA}$)<br>$\alpha$, $\beta$, $\gamma$ ($^\circ$) | 63.31, 108.93, 48.17<br>90, 90, 90 | 64.22, 110.73, 48.56<br>90, 90, 90 |
| Total reflections | 144428 (2254) | 265913 (27723) |
| Unique reflections | 4452 (264) | 20869 (2056) |
| Multiplicity | 32.4 (8.5) | 12.7 (13.5) |
| Completeness (%) | 62.8 (52.9) | 99.47 (99.61) |
| Mean $I/\sigma(I)$ | 6.12 (0.75) | 9.77 (1.41) |
| Wilson *B* factor ($\text{\AA}^2$) | 47.93 | 45.40 |
| $R_{\text{merge}}$ | 0.553 (3.776) | 0.1593 (2.302) |
| $R_{\text{meas}}$ | 0.561 (3.995) | 0.1663 (2.394) |
| $\text{CC}_{1/2}$ | 0.981 (0.597) | 0.997 (0.752) |
| $\text{CC}^*$ | 0.995 (0.860) | 0.999 (0.927) |

Statistics for the highest-resolution shell are shown in parentheses.

Comparing statistics, we can see that resolution, completeness and I/σ(I) are substantially lower for MicroED, whereas Wilson *B* factors are similar. The quality of the data is highlighted by good CC_1/2_ and CC* values for both XRD and MicroED over their respective resolution range (**Table 2**). Values of *R*_*merge*_ and *R*_*meas*_ are better for XRD but such disparities are expected given the multi-crystal approach and effects of dynamical scattering of electrons (multiple elastic scattering events) in MicroED.

### 3.3 Phasing and model refinement

Phases were obtained by MR using three different search models: i) a truncated version of the atomic coordinates of *Mt*R2lox (PDB ID: 3ee4) prepared similarly as for phasing the MicroED data (and used for refining the XRD model); ii) a truncated version of the atomic coordinates of *Gk*R2lox (PDB ID: 4hr0); iii) the unmodified model of *Sa*R2lox solved by MicroED (PDB ID:6qrz). In all three cases, a single well-contrasted solution was obtained in the space group *P*2_1_2_1_2. Quality indicators are very close for the two truncated models prepared from homologous proteins, with a log-likelihood gain (LLG) of approximately 400, and a final translation function Z-score (TFZ) around 21. The solution obtained using the MicroED model shows much better scores, *i*.*e*., a LLG of 1509 and a TFZ of 40 (**Table 3**). This emphasizes the quality of the model provided by MicroED. Moreover, phases for the XRD data can be obtained without prior MR search, solely by running a step of rigid body (RB) refinement with the MicroED model. Following RB, atomic *B* factor refinement against the XRD data cut at the same resolution as MicroED (3.0 Å) leads to already decent *R*_*work*_ and *R*_*free*_ values of 0.3469 and 0.3621, respectively (**Table 3**). This result highlights further the quality of the *Sa*R2lox structure solved by MicroED.

**Table 3.**
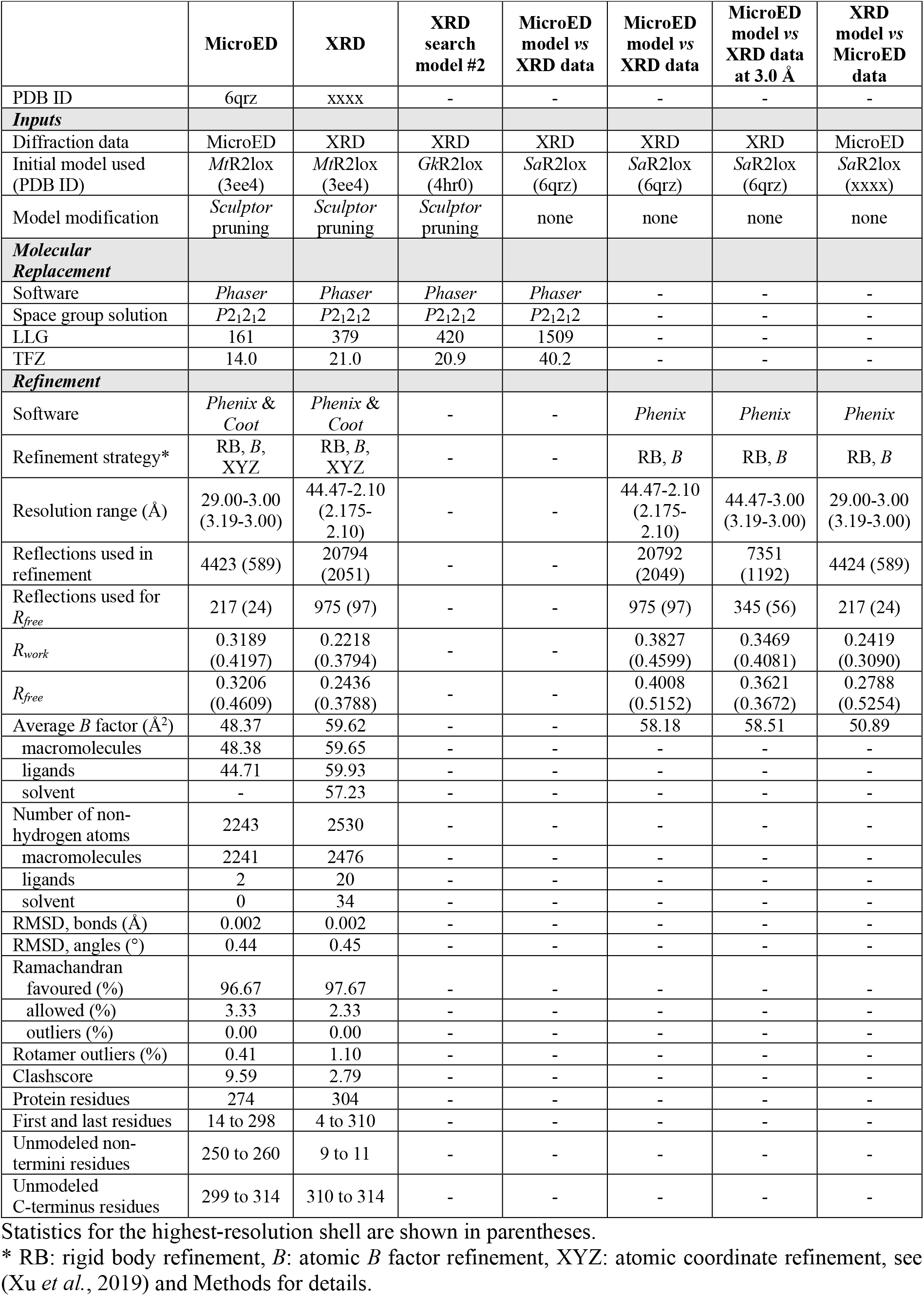
Molecular replacement and refinement statistics of *Sa*R2lox for XRD and MicroED.

|  | MicroED | XRD | XRD search model #2 | MicroED model vs XRD data | MicroED model vs XRD data | MicroED model vs XRD data at 3.0 Å | XRD model vs MicroED data |
| --- | --- | --- | --- | --- | --- | --- | --- |
| PDB ID | 6qrz | xxxx | - | - | - | - | - |
| **Inputs** |  |  |  |  |  |  |  |
| Diffraction data | MicroED | XRD | XRD | XRD | XRD | XRD | MicroED |
| Initial model used (PDB ID) | *MtR2lox* (3ee4) | *MtR2lox* (3ee4) | *GkR2lox* (4hr0) | *SaR2lox* (6qrz) | *SaR2lox* (6qrz) | *SaR2lox* (6qrz) | *SaR2lox* (xxxx) |
| Model modification | *Sculptor* pruning | *Sculptor* pruning | *Sculptor* pruning | none | none | none | none |
| **Molecular Replacement** |  |  |  |  |  |  |  |
| Software | *Phaser* | *Phaser* | *Phaser* | *Phaser* | - | - | - |
| Space group solution | *P2<sub>1</sub>2<sub>1</sub>2* | *P2<sub>1</sub>2<sub>1</sub>2* | *P2<sub>1</sub>2<sub>1</sub>2* | *P2<sub>1</sub>2<sub>1</sub>2* | - | - | - |
| LLG | 161 | 379 | 420 | 1509 | - | - | - |
| TFZ | 14.0 | 21.0 | 20.9 | 40.2 | - | - | - |
| **Refinement** |  |  |  |  |  |  |  |
| Software | *Phenix & Coot* | *Phenix & Coot* | - | - | *Phenix* | *Phenix* | *Phenix* |
| Refinement strategy* | RB, *B*, XYZ | RB, *B*, XYZ | - | - | RB, *B* | RB, *B* | RB, *B* |
| Resolution range (Å) | 29.00-3.00 (3.19-3.00) | 44.47-2.10 (2.175-2.10) | - | - | 44.47-2.10 (2.175-2.10) | 44.47-3.00 (3.19-3.00) | 29.00-3.00 (3.19-3.00) |
| Reflections used in refinement | 4423 (589) | 20794 (2051) | - | - | 20792 (2049) | 7351 (1192) | 4424 (589) |
| Reflections used for $R_{free}$ | 217 (24) | 975 (97) | - | - | 975 (97) | 345 (56) | 217 (24) |
| $R_{work}$ | 0.3189 (0.4197) | 0.2218 (0.3794) | - | - | 0.3827 (0.4599) | 0.3469 (0.4081) | 0.2419 (0.3090) |
| $R_{free}$ | 0.3206 (0.4609) | 0.2436 (0.3788) | - | - | 0.4008 (0.5152) | 0.3621 (0.3672) | 0.2788 (0.5254) |
| Average *B* factor (Å<sup>2</sup>) | 48.37 | 59.62 | - | - | 58.18 | 58.51 | 50.89 |
| macromolecules | 48.38 | 59.65 | - | - | - | - | - |
| ligands | 44.71 | 59.93 | - | - | - | - | - |
| solvent | - | 57.23 | - | - | - | - | - |
| Number of non-hydrogen atoms | 2243 | 2530 | - | - | - | - | - |
| macromolecules | 2241 | 2476 | - | - | - | - | - |
| ligands | 2 | 20 | - | - | - | - | - |
| solvent | 0 | 34 | - | - | - | - | - |
| RMSD, bonds (Å) | 0.002 | 0.002 | - | - | - | - | - |
| RMSD, angles (°) | 0.44 | 0.45 | - | - | - | - | - |
| Ramachandran favoured (%) | 96.67 | 97.67 | - | - | - | - | - |
| allowed (%) | 3.33 | 2.33 | - | - | - | - | - |
| outliers (%) | 0.00 | 0.00 | - | - | - | - | - |
| Rotamer outliers (%) | 0.41 | 1.10 | - | - | - | - | - |
| Clashscore | 9.59 | 2.79 | - | - | - | - | - |
| Protein residues | 274 | 304 | - | - | - | - | - |
| First and last residues | 14 to 298 | 4 to 310 | - | - | - | - | - |
| Unmodeled non-termini residues | 250 to 260 | 9 to 11 | - | - | - | - | - |
| Unmodeled C-terminus residues | 299 to 314 | 310 to 314 | - | - | - | - | - |

Statistics for the highest-resolution shell are shown in parentheses.
\* RB: rigid body refinement, *B*: atomic *B* factor refinement, XYZ: atomic coordinate refinement, see (Xu *et al.*, 2019) and Methods for details.

The final model refined against XRD data was obtained after iterative steps of atomic refinement of coordinates and *B* factors in *phenix*.*refine*, coupled with real-space refinement and model building in *Coot*, performed independently from, but similarly as, for the MicroED structure. Good agreement between the XRD and MicroED models can be observed through superposition of the structures (**Fig. 2*a***). Mean and root mean square (RMS) deviations of Cα for residues 14 to 297 are 0.534 Å and 0.667 Å, respectively (**Fig. 2*b***). Cα deviations are higher in the C-terminus of the protein which moreover includes a 11-residue stretch (250 to 260) unmodeled in the MicroED structure. Note that residue 298, the last residue in the MicroED structure, was excluded from the overall deviation calculations because considered an outlier with a Cα deviation of 6.38 Å compared to the XRD structure, indicating that MicroED did not allow to correctly model it. Besides, refined against the MicroED data, the XRD model gives much better overall *R*_*work*_ and *R*_*free*_ values than the MicroED model (**Table 3**). To note, the large gap between *R* values with a *R*_*free*_ of 0.5254 in the highest resolution shell is expected and indicates logically that the XRD model overfits the lower-resolution MicroED data.

**Figure 2.**
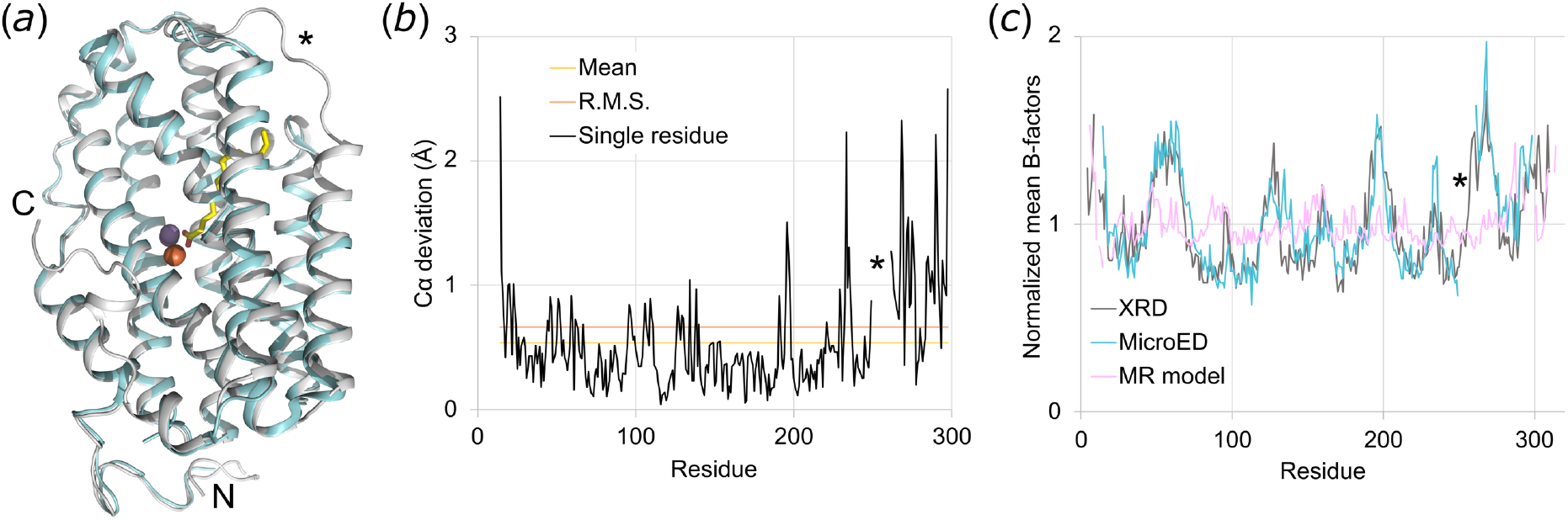
Overall comparison of XRD and MicroED models of *Sa*R2lox. (*a*) Superposition of *Sa*R2lox models obtained by XRD and MicroED (grey and cyan, respectively). Manganese and iron ions are depicted as spheres. The fatty acid ligand is modeled only in the XRD structure. The N- and C-termini are marked N and C, respectively. A star highlights a 11-residue stretch missing from the MicroED structure but modeled in the XRD structure. (*b*) Cα deviation of each residue (from 14 to 297) between the XRD and MicroED structures. The mean and RMS deviations are also indicated (0.534 Å and 0.667 Å, respectively). Note that residue 298 was excluded. (*c*) Normalized mean *B* factors of each residue for the XRD and MicroED structures, and for the search model used for MR.

The quality of the overall geometry of both MicroED and XRD models is comparable, but the average refined *B* factors differ markedly (48.37 and 59.62 Å^2^, respectively), being 23% higher for the XRD model (**Table 3**). However, normalized *B* factors are consistent between MicroED and XRD models across the whole sequence, showing no model-bias from MR and no localized alteration (**Fig. 2*c***). In order to see if such discrepancy could be explained by the modelling of waters and flexible loops in the XRD model, we excluded from the calculation all atoms unmodeled in the MicroED structure. The resulting average refined *B* factor is 58.43 Å^2^, a value still 21% higher than for the MicroED model. Thus, the higher overall value for the XRD structure arises from a global increase of the refined *B* factors, not from differences in molecular structure. Furthermore, this increase is not due to differences in the data processing parameters because the average *B* factor increases from 48.37 to ∼58 Å^2^ when atomic *B* factors of the MicroED model are refined against the XRD data, but decreases from 59.62 to 50.89 Å^2^ when the XRD model is refined against the MicroED data (**Table 3**). A possible explanation for the discrepancy could be the moderate anisotropy of the XRD data, as detected by *phenix*.*xtriage*, which impacts calculation of atomic *B* factors during refinement.

### 3.4. Biologically relevant structural information

The main model improvement obtained by XRD over MicroED is the number of atoms modeled, providing more biologically relevant features for the *Sa*R2lox protein. First, no ligand could be observed in the binding pocket of the protein using MicroED, even though a fatty acid is present and interacts with the metal cofactor as revealed by XRD (**Fig. 3*a***). Moreover, 34 water molecules not visible by MicroED are modeled in the XRD structure, including two coordinating the metal cofactor (**Fig. 3*a***). Additionally, 7 and 12 more residues could be modeled in the N- and C-termini of the protein, respectively (**Table 3**). Finally, a stretch of 11 missing residues in the MicroED structure could be modeled thanks to the better resolved electron density map produced by XRD (**Fig. 3*b***). Besides, it should be mentioned that metal ions could be accurately located in the XRD anomalous difference map (**Fig. 3*a***).

**Figure 3.**
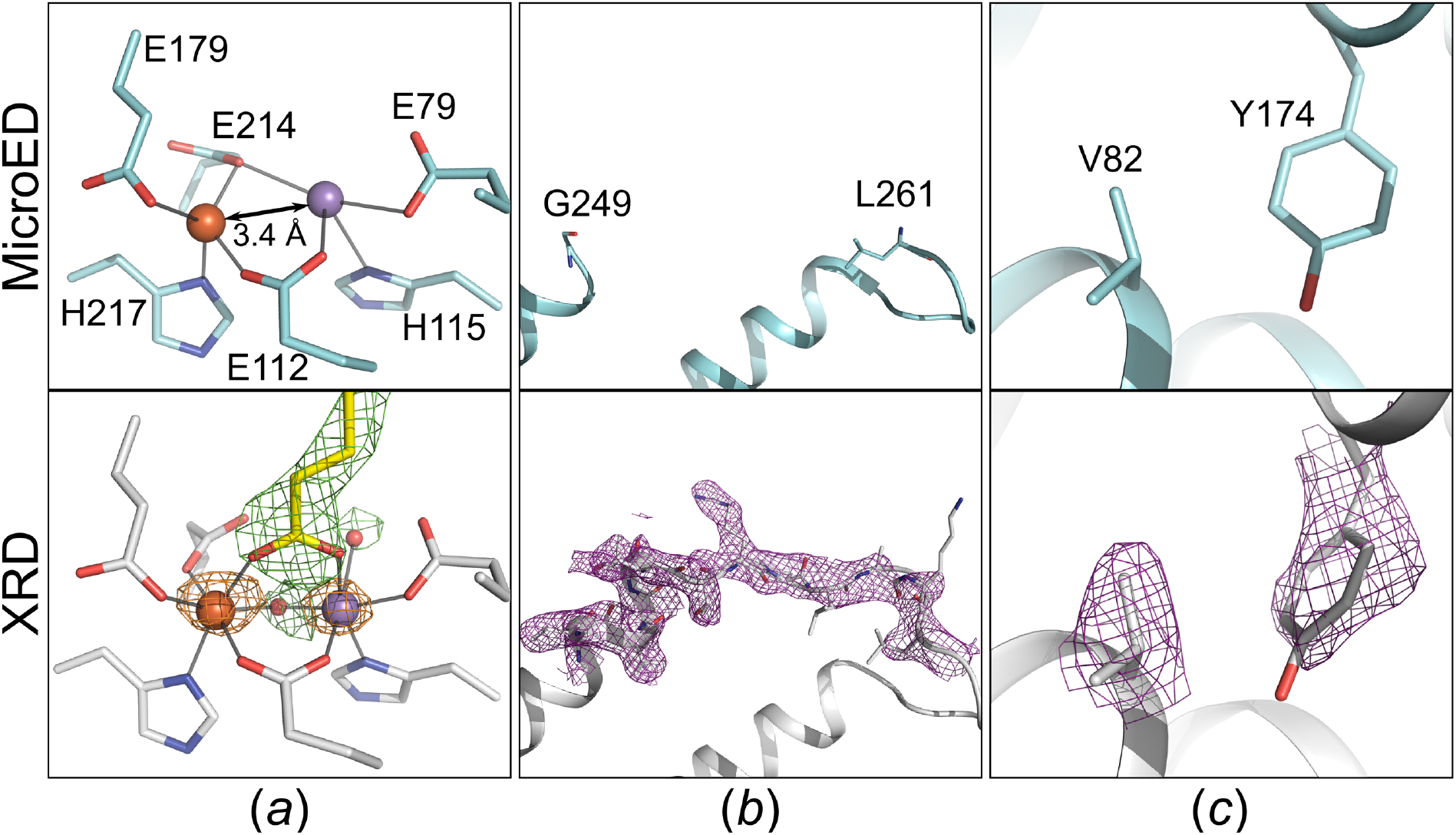
Comparison of biologically relevant features revealed by the XRD and MicroED structures. (*a*) Anomalous difference map (contoured at 5 σ) allows to locate the metal ions, and simulated annealing omit *F*_*obs*_ *– F*_*calc*_ map (contoured at 4 σ) shows the presence of a ligand modeled as palmitic acid and of two metal-coordinating waters in the XRD structure. (*b*) Simulated annealing composite omit maps (contoured at 1 σ) for the 11-residue stretch missing from the MicroED structure but modeled in the XRD structure, and (*c*) the valine and tyrosine residues usually forming an ether cross-link in this protein family but not in *Sa*R2lox.

Despite the lower resolution, MicroED data allowed not only to solve a new protein structure with an α-helix bundle core characteristic of this metalloenzyme family, but also unveiled interesting features specific to *Sa*R2lox such as a distinct substrate-binding pocket (Xu *et al*., 2019). Furthermore, in the two previously known R2lox structures, a tyrosine-valine ether cross-link was present in the protein scaffold. Nevertheless, this cross-link was not modeled in the new *Sa*R2lox structure because the electrostatic potential map obtained by MicroED supported that the two residues were too far apart to form a covalent bond. Importantly, the absence of cross-link is confirmed by the structure redetermined by XRD (**Fig. 3*c***). In addition, even if the coordination sphere of the metal centre was incompletely modeled by MicroED, the distance of 3.4 Å between the metal ions is confirmed in the XRD structure, while the structure used to build the MR search model exhibits a distance of 3.6 Å.

## 4. Discussion

In this work, we redetermined the crystal structure of *Sa*R2lox which is to date the only novel protein structure obtained using MicroED. In light of our studies, we have summarized in **Table 4** the specific challenges encountered and strategies adopted to determine the crystal structure of *Sa*R2lox by MicroED and XRD. This comparison illustrates the intrinsic strengths and weaknesses of the two methods in the case of this real-life protein.

**Table 4.** Challenges encountered and strategies adopted for *Sa*R2lox structure determination by MicroED and XRD.

| Challenges | Strategies |  |
| --- | --- | --- |
|  | MicroED | XRD |
| ***Selection of crystals*** |  |  |
| Visual inspection | Difficult to distinguish microcrystals from precipitate using light microscope; TEM screening of individual crystallization conditions necessary | Use of light microscope |
| Crystallization optimisation | Rapid after identification of the initial hit | Exhaustive screening of conditions around the initial hit (pH, precipitant concentration, drop volume and ratio) |
| Test for diffraction quality | Concomitant to the TEM screening of crystallization conditions | Screening of mounted and cryo-cooled single crystals at synchrotron |
| ***Crystal handling*** |  |  |
| Fragility of crystals | Not a limitation | Repeated manual mounting of individual crystals grown in clusters |
| Opacity of crystallization drops | Not a limitation | Fast crystal mounting |
| Viscous mother liquor | Easily manageable using *Preassis* | Laborious manual mounting |
| ***Data collection & processing*** |  |  |
| Radiation source | Electrons from a standard TEM | Requirement of synchrotron |
| Processing | Use of standard software | Use of standard software |
| ***Biologically relevant insights*** |  |  |
| Metal ion localization | Based on previous studies | Located using anomalous signal |
| Metal ion oxidation state | Based on previous studies but identifiable in theory | Based on previous studies; oxidation states are not identifiable by XRD |
| Presence of a ligand | Not detected | Visible in the electron density map |
| Absence of an ether cross-link | Supported by the electrostatic potential map and correctly modeled | Confirmed by the electron density map |

The first step in our MicroED and XRD studies was to select an optimal crystallization condition. Here, it is necessary to be able i) to identify crystals of suitable size by visual inspection, and ii) to assess their diffraction quality. On one hand, although MicroED requires only little optimization once an initial hit is found, it is difficult to identify nano- and microcrystals as they can fall beyond the resolution limit of light microscopes. Consequently, we had to screen each individual condition using TEM which was very time consuming. This major bottleneck of the method can prevent the identification of suitable crystallization conditions, and therefore call for the development of sample screening solutions. On the other hand, the collection of diffraction data was performed concomitantly to this screening, and therefore did not require any additional step or equipment. In contrast, XRD necessitated an extensive screening of crystallization conditions in order to produce large enough crystals. Moreover, the diffraction quality of these crystals could not easily be tested in parallel, and required the use of a synchrotron.

Regarding crystal harvesting, MicroED appeared to have handy advantages compared to XRD in the case of *Sa*R2lox. Firstly, large crystals for this protein grow in clusters and are very fragile due to their thinness. Secondly, crystallization drops become quickly opaque when exposed outside the well, and thus crystals become rapidly unrecognisable. Thirdly, the mother liquor is particularly viscous due to the high PEG 400 content. These three aspects are commonly encountered in protein crystallography and often render single crystal handling in XRD laborious. By eliminating the need for single crystal manipulation, MicroED greatly simplified the whole process of sample preparation in the case of *Sa*R2lox, especially by using the *Preassis* setup which allows to easily manage viscous crystallization conditions (Zhao *et al*., 2021). Moreover, in such case, MicroED could be not only beneficial from an experimental point of view but also for structure quality. Indeed, unlike MicroED, the XRD structure has substantially higher average refined *B* factor than Wilson *B* factor, possibly because of anisotropy. It is tempting to speculate that the arduous handling of *Sa*R2lox sample in XRD experiment generated excessive strain on the crystal, increasing crystal packing defects and leading to anisotropy. Perturbations apply more uniformly to microcrystals than to larger crystals because of the much smaller volume (Wolff *et al*., 2020). However, further work is needed to know if this systematically impacts refined *B* factors in XRD more than in MicroED.

We showed that the quality of the MicroED data was sufficient to accurately model most of the protein structure and to resolve several important biologically relevant features specific to *Sa*R2lox, such as the absence of an ether cross-link or the metal–metal distance of the protein cofactor. However, XRD data allowed to improve the model and provided further insights into the *Sa*R2lox protein. Importantly, XRD revealed that the metal coordination was incomplete and that the protein ligand had not been modeled in the MicroED structure. Interestingly, even if we did not take advantage of it in the case of *Sa*R2lox, MicroED was shown to allow charge refinement of metal ions in protein models (Blum *et al*., 2021; Yonekura *et al*., 2015). This feature could complement the location-specific information provided by XRD anomalous signal for metal ions. Therefore, MicroED could become a method of choice for the study of metalloproteins.

The lower amount of detail provided by MicroED could likely be imputed to the limited resolution and completeness of the data compared to XRD. These weaknesses are possibly owing to current technical limitations which might be overcome in the future, *e*.*g*., with the development of optimized strategies for sample preparation and data collection, or enhancement of detector sensitivity. However, it is also possible that the gap in resolution limits between XRD and MicroED data is partly due to the different number of unit cells contributing to the diffraction signal, as previously hypothesized (Blum *et al*., 2021). Indeed, taking into account X-ray and electron beam sizes (**Table 2**), diffraction data were produced using crystal volume three to four orders of magnitude larger for XRD than for MicroED. More work is required to carefully investigate this, but such a drawback could weaken the utility of MicroED for protein crystals. Therefore, additional novel protein structures with unique characteristics have to be solved by MicroED in order to confirm the true potential of the method in structural biology.

## Acknowledgements

We thank the staff from the beamline X06SA (PXI) at the Swiss Light Source, Paul Scherrer Institut, Villigen, Switzerland (proposal 20182304), as well as Kristīne Grāve and Hagen Sülzen for their help with data collection.

## Author contributions

H.L. performed protein production, crystallization, data processing, data analysis, project design and wrote the manuscript. H.X., X.Z. and M.H. contributed to project design, data analysis and manuscript writing.

## References

1. Afonine, P. V., Grosse-Kunstleve, R. W., Echols, N., Headd, J. J., Moriarty, N. W., Mustyakimov, M., Terwilliger, T. C., Urzhumtsev, A., Zwart, P. H. & Adams, P. D. (2012). Acta Cryst. D68, 352–367.

2. Andersson, C. S. & Hogbom, M. (2009). Proc. Natl. Acad. Sci. 106, 5633–5638.

3. Blum, T. B., Housset, D., Clabbers, M. T. B., van Genderen, E., Bacia-Verloop, M., Zander, U., McCarthy, A. A., Schoehn, G., Ling, W. L. & Abrahams, J. P. (2021). Acta Cryst. D77, 75–85.

4. Brázda, P., Palatinus, L. & Babor, M. (2019). Science. 364, 667–669.

5. Bunkóczi, G. & Read, R. J. (2011). Acta Cryst. D67, 303–312.

6. Casanas, A., Warshamanage, R., Finke, A. D., Panepucci, E., Olieric, V., Nöll, A., Tampé, R., Brandstetter, S., Förster, A., Mueller, M., Schulze-Briese, C., Bunk, O. & Wang, M. (2016). Acta Cryst. D72, 1036–1048.

7. Clabbers, M. T. B., Fisher, S. Z., Coinçon, M., Zou, X. & Xu, H. (2020). Commun. Biol. 3, 417.

8. Clabbers, M. T. B. & Xu, H. (2020). Drug Discov. Today Technol. S1740674920300354.

9. Clabbers, M. T. B. & Xu, H. (2021). Acta Cryst. D77, 313–324.

10. Emsley, P., Lohkamp, B., Scott, W. G. & Cowtan, K. (2010). Acta Cryst. D66, 486–501.

11. Gemmi, M., Mugnaioli, E., Gorelik, T. E., Kolb, U., Palatinus, L., Boullay, P., Hovmöller, S. & Abrahams, J. P. (2019). ACS Cent. Sci. 5, 1315–1329.

12. Griese, J. J., Roos, K., Cox, N., Shafaat, H. S., Branca, R. M. M., Lehtio, J., Graslund, A., Lubitz, W., Siegbahn, P. E. M. & Hogbom, M. (2013). Proc. Natl. Acad. Sci. 110, 17189–17194.

13. Huang, Z., Willhammar, T. & Zou, X. (2021). Chem. Sci. 12, 1206–1219.

14. Jones, C. G., Martynowycz, M. W., Hattne, J., Fulton, T. J., Stoltz, B. M., Rodriguez, J. A., Nelson, H. M. & Gonen, T. (2018). ACS Cent. Sci. 4, 1587–1592.

15. Kabsch, W. (2010). Acta Cryst. D66, 125–132.

16. Krissinel, E. & Henrick, K. (2004). Acta Cryst. D60, 2256–2268.

17. Kunde, T. & Schmidt, B. M. (2019). Angew. Chem. Int. Ed. 58, 666–668.

18. Liebschner, D., Afonine, P. V., Baker, M. L., Bunkóczi, G., Chen, V. B., Croll, T. I., Hintze, B., Hung, L.-W., Jain, S., McCoy, A. J., Moriarty, N. W., Oeffner, R. D., Poon, B. K., Prisant, M. G., Read, R. J., Richardson, J. S., Richardson, D. C., Sammito, M. D., Sobolev, O. V., Stockwell, D. H., Terwilliger, T. C., Urzhumtsev, A. G., Videau, L. L., Williams, C. J. & Adams, P. D. (2019). Acta Cryst. D75, 861–877.

19. McCoy, A. J., Grosse-Kunstleve, R. W., Adams, P. D., Winn, M. D., Storoni, L. C. & Read, R. J. (2007). J. Appl. Crystallogr. 40, 658–674.

20. Moriarty, N. W., Grosse-Kunstleve, R. W. & Adams, P. D. (2009). Acta Cryst. D65, 1074–1080.

21. Nannenga, B. L. (2020). Struct. Dyn. 7, 014304.

22. Nannenga, B. L. & Gonen, T. (2019). Nat. Methods. 16, 369–379.

23. Williams, C. J., Headd, J. J., Moriarty, N. W., Prisant, M. G., Videau, L. L., Deis, L. N., Verma, V., Keedy, D. A., Hintze, B. J., Chen, V. B., Jain, S., Lewis, S. M., Arendall, W. B., Snoeyink, J., Adams, P. D., Lovell, S. C., Richardson, J. S. & Richardson, D. C. (2018). Protein Sci. 27, 293–315.

24. Wolff, A. M., Young, I. D., Sierra, R. G., Brewster, A. S., Martynowycz, M. W., Nango, E., Sugahara, M., Nakane, T., Ito, K., Aquila, A., Bhowmick, A., Biel, J. T., Carbajo, S., Cohen, A. E., Cortez, S., Gonzalez, A., Hino, T., Im, D., Koralek, J. D., Kubo, M., Lazarou, T. S., Nomura, T., Owada, S., Samelson, A. J., Tanaka, T., Tanaka, R., Thompson, E. M., van den Bedem, H., Woldeyes, R. A., Yumoto, F., Zhao, W., Tono, K., Boutet, S., Iwata, S., Gonen, T., Sauter, N. K., Fraser, J. S. & Thompson, M. C. (2020). IUCrJ. 7, 306–323.

25. Xu, H., Lebrette, H., Clabbers, M. T. B., Zhao, J., Griese, J. J., Zou, X. & Högbom, M. (2019). Sci. Adv. 5, eaax4621.

26. Xu, H., Lebrette, H., Yang, T., Srinivas, V., Hovmöller, S., Högbom, M. & Zou, X. (2018). Structure. 26, 667–675.e3.

27. Yonekura, K., Kato, K., Ogasawara, M., Tomita, M. & Toyoshima, C. (2015). Proc. Natl. Acad. Sci. 112, 3368–3373.

28. Zhao, J., Xu, H., Lebrette, H., Carroni, M., Taberman, H., Högbom, M. & Zou, X. (2021). bioRxiv, https://doi.org/10.1101/665448.

